# Brain-Derived Extracellular Vesicles are Highly Enriched in the Prion Protein and Its C1 Fragment: Relevance for Cellular Uptake and Implications in Stroke

**DOI:** 10.1101/850099

**Authors:** Santra Brenna, Hermann C. Altmeppen, Behnam Mohammadi, Björn Rissiek, Florence Schlink, Peter Ludewig, Antonio Virgilio Failla, Carola Schneider, Markus Glatzel, Berta Puig, Tim Magnus

**Author notes:** Joint senior authorship. To whom correspondence should be addressed.

## Abstract

Extracellular vesicles (EVs) are important means of intercellular communication and a potent tool for regenerative therapy. In ischemic stroke, transient blockage of a brain artery leads to a lack of glucose and oxygen in the affected brain tissue, provoking neuronal death by necrosis in the core of the ischemic region. The fate of neurons in the surrounding penumbra depends on the stimuli, including EVs, received during the following hours. A detailed characterization of such stimuli is crucial not only for understanding stroke pathophysiology but also for new therapeutic interventions.

In the present study, we characterize the EVs in mouse brain under physiological conditions and 24h after induction of transient ischemia in mice. We show that, in steady-state conditions, microglia are the main source of small EVs (sEVs) whereas after ischemia, the main EV population originates from astrocytes. Moreover, sEVs presented high amounts of the prion protein (PrP) which were increased after stroke. Conspicuously, sEVs were particularly enriched in a truncated PrP fragment (PrP-C1). Because of similarities between PrP-C1 and certain viral surface proteins, we studied the cellular uptake of brain-derived sEVs from mice lacking (PrP-KO) or containing PrP (WT). We show that PrP-KO-EVs are rapidly taken up by neurons and colocalize with lysosomes. Although eventually WT-EVs are also found in lysosomes, the amount taken up by neurons is significantly higher for PrP-KO-EVs. Likewise, microglia and astrocytes were also engulfing PrP-KO-sEVs more efficiently than WT-sEVs.

Our results provide information on the relative contribution of brain cell types to the sEV pool in mice and indicate that increased release of sEVs by astrocytes together with elevated levels of PrP in sEVs may play a role in intercellular communication at early stages after stroke. In addition, amounts of PrP (and probably PrP-C1) in brain sEVs seem to contribute to their cellular uptake.

## INTRODUCTION

Extracellular vesicles (EVs) are lipid bilayer structures released from almost all types of cells, that carry biologically active molecules, such as proteins, lipids, and extracellular RNAs (e.g. mRNA, miRNA, tRNA and YRNA), and are capable to elicit responses in the receptor cells ^1, 2^. Although once considered as “platelet dust” or “trash cans”, EVs are currently regarded as potent means of intercellular communication and currently represent an intense field of research ^3–6^. Exosomes (with a size of 40-150 nm, originating from multivesicular bodies), microvesicles (150-1,000 nm, shed from the plasma membrane) and apoptotic vesicles (ApoEVs; 1,000-5,000 nm, released from cells undergoing apoptosis) are subtypes of EVs. An increasing amount of evidence shows that EVs play important roles in physiological and in pathological conditions ^7–9^. Moreover, they are considered as potential disease biomarkers and, given their ability to cross the blood-brain barrier (BBB), are also investigated as tools for tissue- or cell-specific delivery of a therapeutic cargo ^10–14^.

Stroke is the second most common cause of death and the main cause of disability worldwide, being responsible for ∼6 million deaths in 2016 (http://www.who.int/en/news-room/fact-sheets/detail/the-top-10-causes-of-death). Ischemic stroke accounts for 87% of cases and is caused by an occluded brain artery, which leads to a temporary lack of glucose and oxygen supply in the affected brain region. Neurons –the most susceptible brain cell population– that are at the core of the stroke will die by necrosis, whereas neurons located at the periphery (penumbra) will enter into electrical silence and, depending on many factors, will either die or survive ^15, 16^. The pathophysiology of stroke is very complex and involves several mechanisms such as excitotoxicity and neuroinflammation ^17–19^. At present, the only therapeutic approach is recanalization and treatment with recombinant tissue plasminogen activator (rtPA). The latter has a very limited time window of 4.5 to 6 hours after stroke and, thus, only 20% of the stroke patients can benefit from it ^20^. Hence, there is an urgent need for novel therapeutic options to be used after this time period. At present, EVs are regarded as therapeutic tools for regenerative therapy after stroke ^21, 22^.

The cellular prion protein (PrP) is enriched in EVs isolated from cerebral spinal fluid (CSF) and in neuronal cells ^23, 24^. PrP is a cell surface N-glycosylated, GPI-anchored protein present in detergent-resistant domains (also known as lipid rafts) ^25^, which are important for the biogenesis of EVs ^26, 27^. Although PrP is highly conserved through evolution and many functions have been suggested, the exact physiological role of PrP is still not well-defined, mainly because knock-out mice for PrP do not present major deficiencies ^28, 29^. However, under ischemic conditions, it has been shown that PrP has a protective role. Thus, in mouse models of stroke, PrP^C^-deficient (PrP-KO) mice present an increased stroke volume compared to wild-type (WT) mice ^29, 30^, which can be rescued by PrP overexpression ^31, 32^. Moreover, mice overexpressing PrP^C^ showed improved long-term neuronal recovery after stroke, which was associated with increased neuro- and angiogenesis ^31^. Interestingly, in mouse models of ischemia, but also in brains from patients suffering from stroke, an increase of PrP has been observed at the penumbra area, probably as an attempt to decrease the oxidative stress ^33–, 35^. Last but not least, exosomal PrP secreted by astrocytes under ischemic conditions, had a protective effect on ischemic neuronal cerebellar cells, but this protection was eliminated when exosomes were exempt from PrP ^36^. PrP undergoes several physiological cleavages that are highly conserved in mammals, including the α- and β-cleavage, and shedding near the GPI anchor, which may account for the myriad of functions attributed to PrP ^37–39^. The fact that α-cleavage, leading to the formation of a released neuroprotective N1 fragment and a membrane-attached C1 fragment, seems to be performed and ensured by several (yet-to-be-identified) proteases, provides an idea of its biological importance ^37, 40^.

Because of the potential of EVs in stroke therapy ^21, 41^, further understanding and characterization of brain-derived EVs in physiological and under stroke conditions becomes necessary. With this aim, and taking advantage of the recently developed protocols to isolate EVs from brain ^42^, we show here that (i) microglia are the main cell population to release small EVs (≤ 200nm, sEVs) in steady-state conditions; (ii) brain-derived sEVs are enriched in PrP and especially in its truncated C1 fragment, and (iii) this influences the cellular uptake of EVs. Moreover, (iv) lack of PrP on sEVs increases their uptake by neurons, microglia, and astrocytes. Finally, (v) in a mouse model of stroke, astrocytic contribution to the sEV pool as well as levels of PrP and PrP-C1 on brain sEVs are significantly increased after 24h of reperfusion. We hypothesize that regulating the amounts of PrP, and particularly PrP-C1, is a mechanism to modulate EVs uptake by brain cells and that a deeper understanding of the increased astrocytic release of EVs in reaction to stroke could lead to a targeted therapy.

## MATERIAL AND METHODS

### Ethics statement

All animal experiments have been conducted after the approval of the local animal care committees (*Behörde für Lebensmittelsicherheit und Veterinärwesen Hamburg*, project number: N45/2018) and in accordance with the guidelines of the animal facility of the University Medical Center Hamburg-Eppendorf.

### Transient Middle Cerebral Artery Occlusion (tMCAO)

The tMCAO was performed as previously described in detail ^19^. Three-month-old male mice were used for the procedure. tMCAO was achieved by using a 6-0 nylon monofilament to stop the blood supply for 40 minutes. In the control group (“sham”), animals were also anesthetized, and the arteries were visualized but not disturbed.

### Isolation and purification of brain EVs

EVs were isolated from brain as previously described ^42^ with some modifications. Briefly, frozen brains from C57BL/6 WT or *Prnp*^0/0^ (PrP-KO) mice ^43^ were gently chopped in few drops of Hibernate-E (Gibco) and transferred to a 15 mL tube containing 75 U/ml of collagenase type III (Worthington) in Hibernate-E at a ratio of 800 μL buffer per 100 mg of brain, and incubated in a water bath at 37°C for 20 min. Alternatively, some samples were incubated with 75U/mL of collagenase type IV (Worthington). During this time, the tube was mixed by inversion every 5 min and pipetted up and down using a 10 mL pipette. Immediately after, the tube was returned on ice and proteinase inhibitors (cOmplete Protease Inhibitor Tablets, Roche) were added. The sample was then centrifuged at 300x*g* for 5 min at 4°C and the supernatant collected and further centrifuged at 2,000x*g* for 10 min at 4°C. The supernatant was again collected and centrifuged at 10,000x*g* for 30 min at 4°C. In order to isolate small extracellular vesicles (sEVs), the supernatant was passed through a 0.22 µm sodium acetate filter (GE Healthcare). The resulting flow-through (filtered) or supernatants that did not pass through the filter (non-filtered) was layered on top of a sucrose gradient (0.6M, 1.3M, 2.5M). The gradient was centrifuged at 180,000x*g* for 3 h at 4°C and six fractions of 2 mL each were collected, diluted in PBS, and further centrifuged at 100,000xg at 4°C for 1 h. The final pellet was then resuspended in PBS or RIPA buffer (50mM Tris-HCl pH=7.4, 150mM NaCl, 1% NP40, 0.5% Na-Deoxycholate and 0.1% SDS), containing protease and phosphatase inhibitors (Phospho-Stop Tablets, Roche).

### Nanoparticle tracking analysis (NTA)

Resuspended pellets resulting from the 100,000x*g* centrifugation step were used for NTA. Briefly, 1 µL of the final pellet suspension were diluted at 1:5,000 for non-filtered EVs and 1:1,000 for filtered EVs in PBS and 500 µL were loaded into the sample chamber of an LM10 unit (Nanosight, Amesbury, UK). Ten videos of ten seconds were recorded for each sample. Data analysis was performed with NTA 3.0 software (Nanosight). Software settings for analysis were as follows: detection threshold = 6, screen gain = 10.

### Fluorescence labelling of exosomes

EVs, in a concentration of 10^12^/ml, were incubated with 8µM of FM-143 (Invitrogen) or with 5nM of mCLING (Synaptic Systems) on ice and in the dark for 5 min. After incubation, the reaction was stopped by adding 1% BSA in PBS. The unincorporated dye was then removed from labelled EVs by centrifugation at 100,000x*g* at 4°C for 70 min. The final pellet was resuspended in PBS and immediately used for the experiments.

### Primary neuronal culture

Primary hippocampal neurons were prepared at P0-P2 as previously described ^44^. Briefly, animals were decapitated, the brain was rapidly excised and cleaned from meninges and choroid plexus. Hippocampi were isolated and digested for 30 min at 37°C in 10mM glucose (500 µL/brain) containing 25U of papain (Sigma) and 20 µg/mL DNAse (Sigma). Cells were washed several times with plating medium (MEM, 10mM glucose and 10% horse serum) and, after the last wash, cells were mechanically dissociated. 100,000 cells were plated in 12-well plates containing plating medium and poly-L-lysine-coated 13 mm diameter coverslips. After 5 h, the media was changed to Neurobasal-A medium (Gibco) supplemented with 1% B27 (Thermo Fisher), 0.5% Glutamax (Gibco) and 0.1 % Gentamicin (Gibco). After 24h, arabinoside-C (ARA-C, Tocris) was added in a concentration of 15µM to prevent mitotic, non-neuronal cell proliferation. Neuronal cultures were kept at 37 °C in 5% CO2 for 14 days and half of the medium was replaced every 3 days.

### Co-culturing of neurons with an astrocytic feeder layer

For the co-culture, an astrocyte feeder layer was prepared 21 days in advance from P0/P1 mouse pups. Briefly, after isolating the brain and cleaning meninges and choroid plexus, the cortex was removed and put in GGM (Glial Growth Medium: DMEM supplemented with 1.35% glucose, 1% pen/strep and 10% FCS). After addition of pre-warmed 0.25% trypsin, the tissue was digested for 15 min at 37° C in a water bath while shaking (300 rpm). After that, 50 µg/mL of DNAse I was added for 1 minute and the enzymatic reaction was stopped by adding 4 volumes of GMM. Cells were then washed twice with GGM by centrifuging 5 min at 180x*g*, the cell suspension was passed through a 70 µm cell strainer and the final cell resuspension was plated in T75 flasks with GMM.

For neuronal preparation, P0 mice pups were used as previously described ^45^. Briefly, after extracting the brain and cleaning of meninges and choroid plexus, the isolated hippocampi were digested at 37°C for 15 min in dissection media (1x HBSS, 1% pen-strep, 10mM HEPES and 0.6% glucose) containing pre-warmed 0.25% trypsin. After the enzymatic digestion, DNAse I (20 µg/mL) was added and incubated at RT for 1 min. GMM was added to quench the enzymatic reaction. Cells were then centrifuged for 5 min at 180x*g*. The resulting pellet was gently triturated in Neuronal Maintenance Medium (NMM, Neurobasal medium containing 1% glutamax, 2% B27 and 1% pen-strep) using a 5 mL pipette followed by a 1 mL pipette. Cells were then centrifuged again for 5 min at 180x*g*. The pellet was resuspended in NMM and passed through a 70 µm cell strainer. About 40,000 cells were plated in 24-well plates containing pre-coated poly-L-lysin coverslips and NMM. After 2 h, the coverslips were inverted on the top of the pre-cultured feeder layer with a wax-dot spacer in between. After 24 h, the mitotic inhibitor FUDR (2′-Deoxy-5-fluorouridine) was added in a concentration of 10µM. Neuronal cultures were kept at 37°C in 5% CO_2_ for 14 days and half of the medium was replaced every 3 days.

### Mixed glia culture

Mixed glia cultures were prepared from P0-P2 mice pups. Briefly, animals were decapitated, the brain excised and cleaned from meninges. The cortex was isolated and washed twice with HBSS (Gibco) containing 10mM HEPES by centrifuging at 310x*g* for 5 min at 4°C. The tissue was then incubated in digestion solution (HBSS-HEPES 10mM containing 25U of papain and 10 µg/mL DNAse) at 37°C for 30 min. Plating media (BME with 10% FCS and 0.1% gentamycin (Gibco)) was added to stop the digestion reaction. Cells were then further centrifuged 5 min at 310x*g*, the pellet resuspended in the plating media using a Pasteur pipette and passed through a 70 µm strainer. Cells were then plated in T25 Flasks or in 12-well plates.

### Immunocytochemistry, confocal microscopy, and STED

Neurons were incubated for 1, 3 or 6 h with 5 µL (containing about 2.6*10^8^ particles as measured with NTA) of EVs isolated either from WT or PrP-KO brains, previously labelled with mCLING as described above. For the co-culture system, coverslips containing neurons were transferred to another well where the incubation with EVs was taking place without the astrocytic feeder layer. After incubation, neurons were fixed with 4% PFA and 0.2% glutaraldehyde in PBS for 10 min, permeabilized for 10 min with 0.5% saponin in PBS, blocked for 30 min with 1% BSA in PBS-Tween (0.1%) and then incubated for 1 h with the primary antibodies (the lysosomal marker LAMP-1 (Invitrogen, 4-1071-82, 1:500) or the microglia marker IBA1 (Wako, 019-19741, 1:500)) diluted in PBS-0.1% BSA. The coverslips were then washed 3 times with PBS and incubated with the secondary antibody Alexa Fluor donkey anti-rat 488 (Life Technologies, A21208), Alexa Fluor donkey anti-rabbit 555 (Life Technologies, A31572) and Alexa Fluor donkey anti-rat 555 (Abcam, ab150154, 1:500). Actin labelling was performed by adding at this step Phalloidin-iFluor 488 (Abcam, ab176753, 1:500) diluted in BSA 0.1% in PBS. The coverslips were washed 3 times with PBS and mounted with DAPI Fluoromount-G (SouthernBiotech, 0100-20). Between 15-20 pictures of randomly chosen single neurons per experiment were taken with the 63X immersion oil objective and at a magnification of 3X in a Leica TCS SP8 confocal microscope. The signal of the 633 nm wavelength corresponding to EVs was quantified and referred to the cell area using ImageJ. Experiments were repeated 3 times for each time point.

For STED microscopy, a drop of sEVs, labelled with mCLING as previously described, was placed on a coverslip and then mounted with solid mounting medium (Abberior).

STED and corresponding confocal microscopy were carried out in sequential line scanning mode using an Abberior STED expert line microscope. This setup was based on a Nikon Ti-E microscope body and employed for excitation and detection of the fluorescence signal a 60x (NA 1.4) P-Apo oil immersion objective. One pulsed laser was used for excitation at 640 nm and near-infrared pulsed laser (775 nm) for depletion. The detected fluorescence signal was directed through a variable-sized pinhole (set to match 1 Airy at 640 nm) and detected by novel state of the art avalanche photo diodes APDs with appropriate filter settings Cy5 (615-755 nm). Images were recorded with a dwell time of 3 µs and the pixel size was set to be 20 nm. The acquisitions were carried out in time gating mode i.e. with a time gating width of 8 ns and a delay of 781 ps. After acquisition images were displayed and analysed by the freeware ImageJ Fiji.

### Flow cytometry

Mixed culture glia cells were incubated with 5 µL (containing about 2.6*10^8^ particles as measured with NTA) of WT or PrP-KO EVs labeled with mCLING for 3 h. Cells were then trypsinized and transferred to FACS tubes containing FACS buffer (PBS with 1mM EDTA and 0.2% BSA). Cells were centrifuged for 5 min at 4°C at 310x*g*, washed in FACS buffer and centrifuged again. Cells were stained for 30 min on ice with anti-CD11b–FITC (1:100, Clone M 1/70, Biolegend) and anti-GLAST-PE (1:100, Miltenyi Biotec) in the presence of Fc Block (1:100, Bio X Cell) in FACS buffer, washed 2 times with FACS buffer and finally resuspended in 200 µL of FACS buffer. Measurements were done with BD FACSCanto™ II and analyzed with FlowJo.

### Western blot

The protein amount resulting from the 100.000x*g* pellet resuspended in RIPA buffer was determined using the Pierce BCA Protein Assay Kit (Thermo Scientific). Samples were then denatured at 70°C for 10 min with NuPage LDS Sample Buffer (Invitrogen) and NuPage sample reducing agent (Invitrogen) and equal amounts of protein were then loaded on precast NuPage 10% Bis-Tris protein gels (Invitrogen). After electrophoretic separation, proteins were transferred onto nitrocellulose membranes (LI-COR) by wet-blotting. The membranes were then stained with Revert Total Protein Stain (LI-COR), following the manufactureŕs protocol, to detect total amounts of protein. Membranes were subsequently blocked for 1 h with Rotiblock (Roth) and incubated with primary antibody overnight at 4°C on a shaking platform. The antibodies used were: rabbit antibody against Alix (1:500; ABC40, Millipore), mouse monoclonal against CNP (1:1000; C5922, Sigma), rabbit monoclonal against CD81 (1:1000; 10037, Cell Signalling), rabbit antibody against EAAT1 (1:1000; NB100-1869SS, Novusbio); rabbit antibody against EAAT2 (1:500; NBP1-20136SS, Novusbio); mouse antibody against flotillin-1 (1:1000; 610820, BD Biosciences), mouse antibody against GM130 (1:1000; 610822, BD Biosciences), rabbit antibody against PLP (1:1000; NB100-74503, Novusbio), rabbit antibody against P2Y12 (1:500; LS-C209714, LSBio), mouse monoclonal antibodies against PrP (POM1 1:2000; and for some experiments also POM2 at the same dilution ^46^), rabbit antibody against SNAP25 (1:1000; 3926, Cell Signalling); rabbit antibody against synapsin1 (1:1000; 106103, Synaptic Systems) and mouse monoclonal antibody against TMEM119 (1:1000; 66948-1-Ig, Proteintech). After washing with TBST, membranes were incubated for 1 h with the respective HRP-conjugated secondary antibodies (Cell Signalling, 1:1000) and subsequently washed 6× with TBST. After incubation with Pierce ECL Pico or Super Signal West Femto substrate (Thermo Fisher Scientific), chemiluminescence was detected with a ChemiDoc imaging station (BioRad) and quantified using Image Studio software (LI-COR).

### PNGase F assay

For removal of N-linked glycans attached to PrP and its C1 fragment, sEV samples and total homogenates (TH) were digested with PNGase F (New England Biolabs) according to the manufactureŕs protocol.

### Electron microscopy

Pellets from the 100.000x*g* centrifugation were fixed with 4% PFA containing 2.5% glutaraldehyde in PBS and adsorbed in cellulose capillary tubes. Subsequently, the pellets were washed with PBS, post-fixed for 30 minutes with 1% OsO4 in PBS, washed with ddH2O, and stained with 1% uranyl acetate in water. The samples were gradually dehydrated with ethanol and embedded in Epon resin (Carl Roth, Germany) for sectioning. Ultrathin 50 nm sections were prepared using Ultracut Microtome (Leica Microsystems, Germany). The sections were poststained with 2% uranyl acetate. Electron micrographs were obtained with a 2K wide-angle CCD camera (Veleta, Olympus Soft Imaging Solutions GmbH, Münster, Germany) attached to a FEI Tecnai G 20 Twin transmission electron microscope (FEI, Eindhoven, The Netherlands) at 80kv.

### Statistical analysis

The GraphPad Prism 8 statistic software program was used for the statistical analysis. To assess differences between the THs and the corresponding sEVs and between sEVs from sham and strokes in western blot, after passing a normality test, either a paired or unpaired *t*-test was used, respectively. Nanosight measurements were assessed by unpaired *t*-test. Statistical significance was considered when *p*-values were as follows: \**p* < 0.05, \*\**p* < 0.005, and \*\*\**p* < 0.001. Values are given as mean ± SEM. The exact *p*-value is given in the text.

## RESULTS

### Characterization of brain-derived EVs

Recently, the isolation of EVs from brain has become an important tool to study crucial events in the propagation of misfolded proteins in neurodegenerative diseases ^47^. We took advantage of a recently published protocol ^42^ to study the expression of PrP in brain-derived EVs and to assess the cell populations that mainly releases EVs in both, steady-state conditions and after stroke, by using the tMCAO mouse model. Since the new guidelines of the *International Society for Extracellular Vesicles* recommend to differentiate between small EVs (sEVs, ≤200nm) and medium/large EVs (≥ 200nm) instead of exosomes and microvesicles/ectosomes ^48^, we introduced in the above-mentioned protocol an extra filtration step with a 0.22 µm filter before the samples were loaded in the sucrose gradient to differentiate between these two EVs pools (see Material and Methods). As shown in Fig. 1A, six fractions either from non-filtered or filtered samples were collected and analysed by western blot. In the non-filtered samples, a higher amount of total protein (as shown with the total staining, TS) was detected in all fractions compared to the filtered samples, but in both cases fractions 3 to 5 (corresponding to a density of 1.08 to 1.23 g/mL as measured with a refractometer) were enriched in known markers of EVs such as the cytosolic protein Alix (96 kDa), the membrane-bound flotillin-1 (49 kDa), and the tetraspanin CD81 (26 kDa) ^48, 49^. GM130 (130 kDa), a resident protein of the cis-Golgi, was used as “proof of no contamination” marker. The total brain homogenate (TH) is shown for comparison. Furthermore, fractions 3 and 4 were pooled and analyzed with transmission electron microscopy (TEM). Non-filtered samples show a combination of small and large vesicles, whereas in filtered samples the population appears more homogeneous (Fig. 1B). To confirm that the EVs could be visualized by microscopy for further experiments, we labelled them using the mCLING dye, which has the capacity to emit fluorescence only when intercalated in the lipid bilayer but not in the aqueous medium^50^. When isolated EVs were stained, fixed and mounted on coverslips, we compared the signal obtained with confocal microscopy to the one obtained with *Stimulated Emission Depletion* (STED) microscopy (Fig. 1C). Due to the higher resolution, the blurred signal obtained with confocal microscopy (in green) was transformed to single dots corresponding to individual EVs in STED (in red), although some aggregates (probably corresponding to clustered EVs) were also present.

**Fig. 1.**
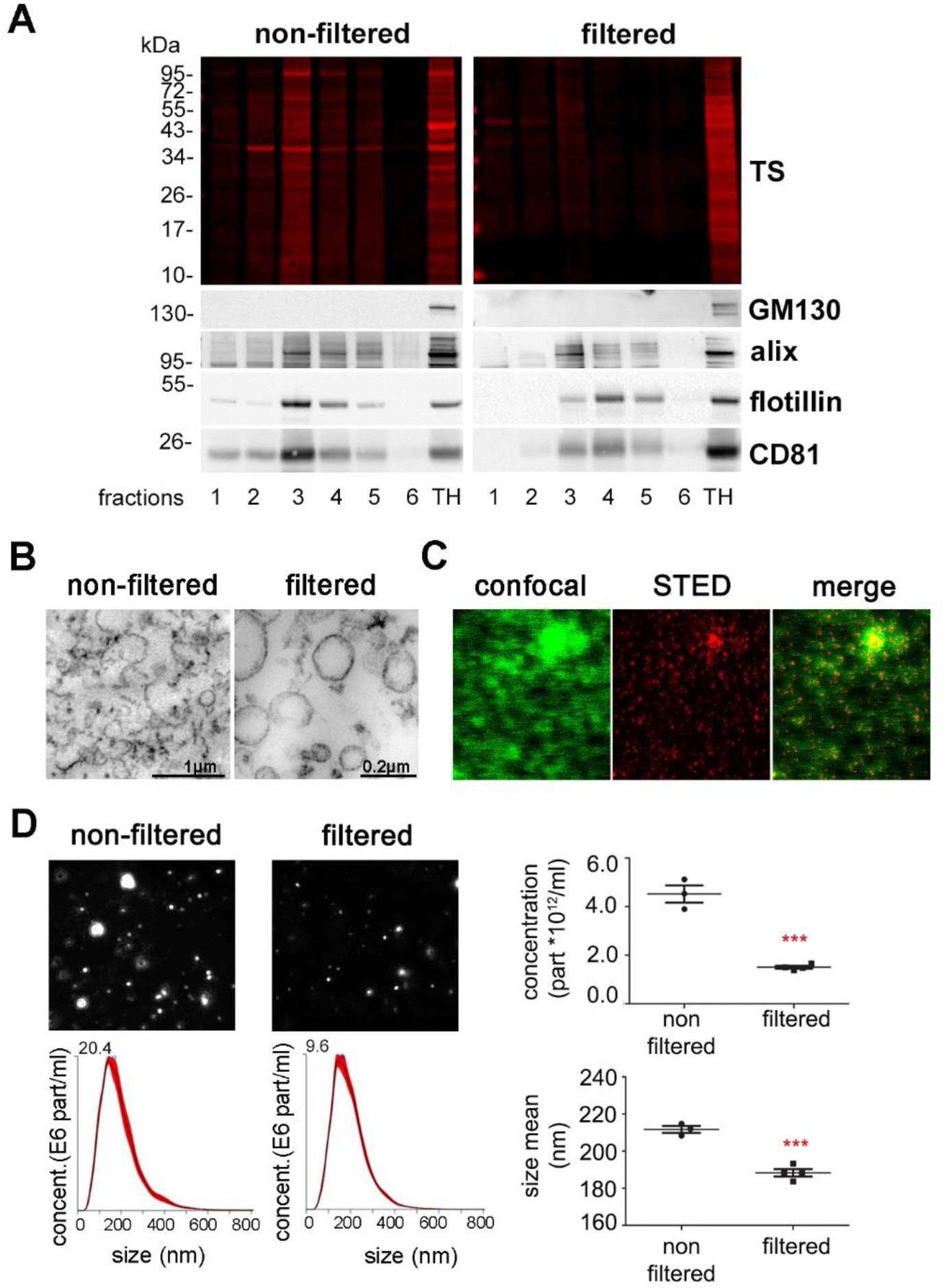
Characterization of brain EVs. (A) Representative western blots of the six fractions obtained after sucrose gradient centrifugation, either from samples that were passed through a 0.22µm filter (filtered, sEVs) or not (non-filtered EVs). The total protein staining (TS, used as a loading control) shows a decrease in the total protein amount for the filtered samples. The cytosolic protein Alix (96 kDa), the membrane-bound Flotillin-1 (49 kDa) and membrane-bound CD81 (26 kDa), all considered as markers of EVs, were found in both cases mainly in fractions 3 and 4. The Golgi resident protein GM130 was used as a marker of non-EVs (negative control). (B) Representative transmission electron microscopy (TEM) images of pooled fractions 3 and 4 from non-filtered and filtered EVs. As seen for the TS in the western blot, the filtered fraction shows a decrease in EVs and a more uniform population. Scale bare is 1 µm for non-filtered and 0.2 µm for filtered EVs.(C) Confocal (green) and STED (red) images from pooled sEVs labelled with mCLING and the merge of both showing that with high-resolution STED imaging, the blurry dots in the confocal correspond to single (or groups) of particles. (D) Representative frames and concentration/size graphs from Nanoparticle Tracking Analysis (NTA) of pooled non-filtered and filtered EVs. Concentration (in particle/mL) and mean size (in nm) analysis of non-filtered pooled EVs (n=3) and filtered pooled EVs (n=4) show a significant reduction of concentration and mean size of EVs after filtration. Values are reported in the main text.

Finally, *Nanoparticle Tracking Analysis* (NTA, Fig. 1D) of both pools revealed a decrease in the number of events counted in the filtered (1.5×10^12^ ±5.8×10^10^) compared to the non-filtered samples (4.5×10^12^ ±3.5×10^11^ *p*= 0.0002), together with a confirmatory decrease in mean EVs size (211.7 ±1.9 nm in non-filtered vs 188.3 ±2 nm in filtered samples *p*= 0.0004). Note that the filtered samples still contain some vesicles that are larger than 200 nm, probably indicating that some large EVs can squeeze through the 0.2 µm filter or representing clustered sEVs

### Assessment of the relative contribution of major cell types to the sEV pool indicates microglia as a dominant source of sEVs in brain under physiological conditions

In order to study the relative contribution of different brain cell populations to the whole pool of brain sEVs in physiological conditions, we assessed enrichment of known brain cell type-specific markers in sEVs relative to their signal in TH by western blot analysis (Fig. 2A). Each sample signal of the western blot was first referred to the respective signal of the TS as a loading control and then, the mean value obtained in sEVs (n=4) was compared to the mean value obtained in the corresponding TH (n=4). Since it could be that a given marker protein is more sorted into sEVs than others (and thus not directly indicating the relative contribution of this cell population to the total pool of sEVs), we assessed two (exclusively membrane-bound) protein markers for each cell population in order to reduce the risk of misinterpretation. The G-protein coupled PY2 receptor (P2Y12) and the Transmembrane protein 119 (TMEM119) were chosen as microglial markers ^51, 52^; PLP (proteolipid protein, a major component of the myelin sheet) and 2′-3′-Cyclic nucleotide 3′-phosphodiesterase (CNP) were used as oligodendrocyte markers ^53, 54^; synapsin 1 and the Synaptosomal Nerve-Associated Protein 25 (SNAP25), present at the membrane of synaptic vesicles and at the pre/post-synaptic membrane respectively, were assessed as neuronal markers ^55, 56^, and the Excitatory Amino Acid Transporters 1 and 2 (EAAT-1 and EAAT-2) were used as astrocytic markers ^57^. Quantifications are shown in Fig. 2B. We observed that, in steady-state conditions, markers for microglia were mainly contributing to the pool of brain sEVs, as P2Y12 (2.5-fold increased; TH set to 100% ±3.9%; sEVs: 247.5 ±12.4%; *p*=0.0073) and TMEM119 (1.7-fold increased; TH set to 100% ±5.5%; sEVs: 168.7 ±8.6%; *p*=0.0119) were most drastically enriched in the sEVs fractions compared to the TH. Interestingly, TMEM119 presented a band at around 55 kDa in the TH, whereas in the sEV-enriched fractions, a main band was observed around 20 kDa. Because four isoforms have been described for TMEM119, this lower band could correspond either to a truncated version of TMEM119 or to one of the four isoforms, probably also lacking the O-glycan modification ^58^. The oligodendrocyte marker PLP showed a slight yet non-significant increase (TH was set to 100% ±4.7%; sEVs: 138.9% ±11%; *p*=0.1), whereas the other oligodendrocyte marker CNP1 revealed no differences (TH set to 100% ±15.3%; sEVs: 97.2% ±12.4%). In contrast, the neuronal markers synapsin 1 (syn-1; TH set to 100% ±12.9%; sEVs: 78.1% ±4.9%; *p*=0.050) and SNAP-25 (TH set to 100% ±3.7%; sEVs: 22.4% ±4.3%; *p*= 0.0018) were significantly decreased in the sEVs in comparison to the TH, thus implying that neuronal sEVs rather represent a minority in the brain sEV pool under normal conditions. Lastly, the astrocytic markers EAAT-1 and EAAT-2 showed no differences in sEVs compared to the TH (for EAAT-1: TH set to 100% ±4.6%; sEVs: 95.9% ±6.7%/ for EAAT-2: TH set to 100% ±14.8%; sEVs: 48% ±14.8%), indicating presence but no relative enrichment in the total pool (Fig. 2B). Thus, neurons and astrocytes only show a moderate contribution to the total sEVs pool in brain, whereas oligodendrocytes and, significantly, microglial sEVS are relatively enriched.

**Fig. 2.**
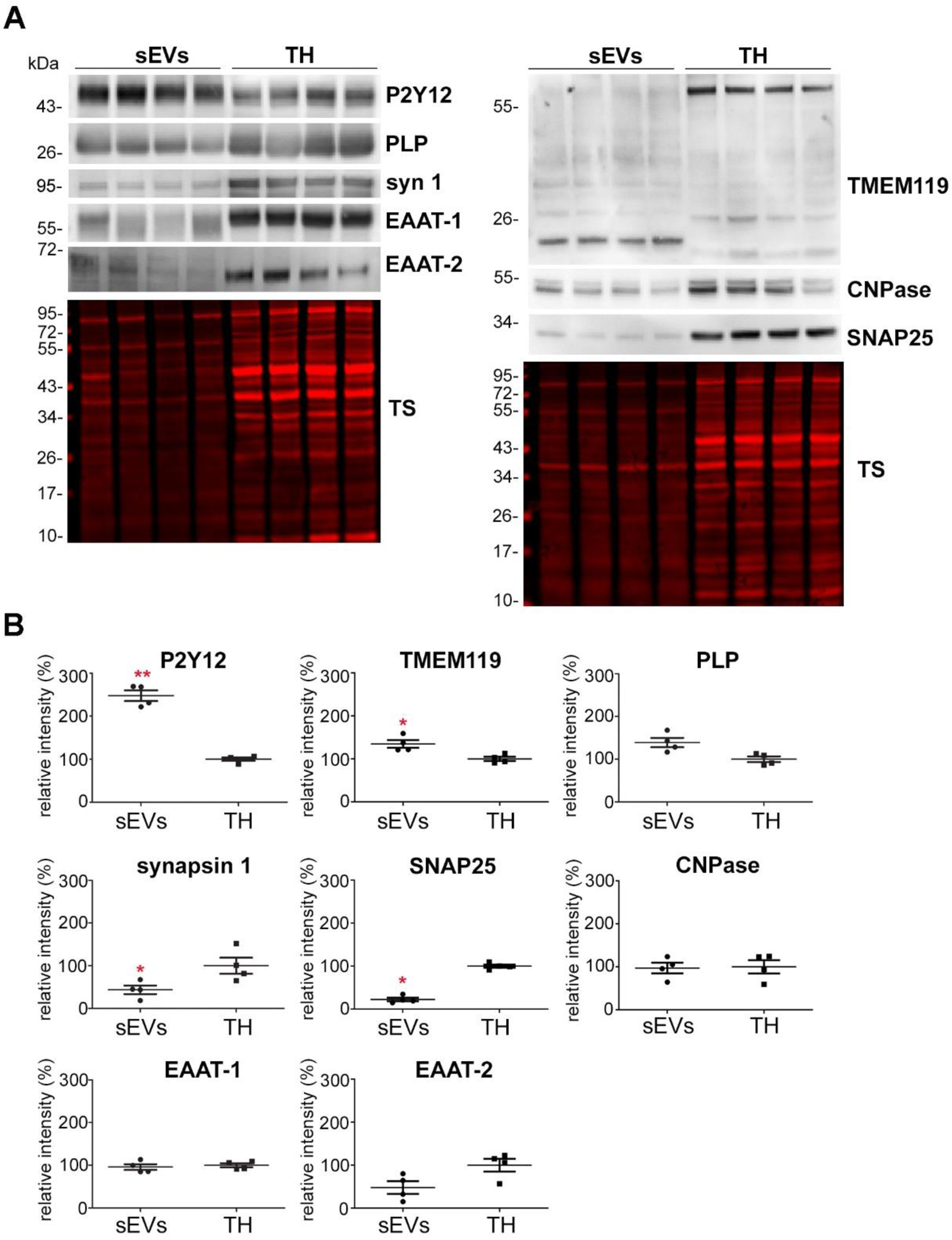
Microglia is the main contributor to the physiological brain sEV pool. (A) Representative western blots of pooled fractions 3 and 4 of filtered EVs (sEVs; n=4) compared to their respective total brain homogenates (TH). P2Y12 and TMEM119 were chosen as markers of microglia, PLP and CNP as markers of oligodendrocytes, synapsin1, and SNAP25 as neuronal markers, and EAAT1 (GLAST) and EAAT2 (GLT-1) as markers of astrocytes. Of note, TMEM119 presented a lower band (at around 20 kDa) in the sEVs-enriched fractions instead of the reported 60 kDa band (approx.) observed in the TH. This could correspond to an isoform or a truncated version of TMEM119. (B) Scatter plots showing relative intensity quantifications of each cell type marker. Each lane was first referred to its total protein staining (TS) and the mean of each group (sEVs vs. TH) was compared in order to check for relative enrichment. P2Y12 and TMEM119 are approximately 2.5 and 1.5 times enriched in sEVS compared to the TH, indicating a dominant contribution from microglia to the whole pool of brain sEVs. Synapsin1 and SNAP25 are significantly decreased compared to the TH, suggesting low contribution of neuronal sEVs to the total pool. PLP, CNP, EAAT1 and EAAT2 did not show any significant differences compared to the TH. Exact mean, SEM and *p* values are given in the main text.

### Brain-derived EVs are enriched in PrP and, in particular, its proteolytically truncated C1 fragment

Since we were also interested in PrP as a protein that was shown to act neuroprotective in ischemic insults, and as a known resident of EVs, we performed western blot analyses of filtered and non-filtered EV-enriched fractions. As shown in Fig. 3A, PrP is present in both, medium/large EVs and sEVs. Interestingly, in both cases, the pattern of PrP in EVs is different compared to the TH when visualized with the POM1 antibody that recognizes the C-terminal part of PrP ^46^. Hence, for the TH we could observe the typical pattern of PrP with a prominent diglycosylated band at around 43 kDa followed by two lower and less conspicuous bands corresponding to mono- and unglycosylated full-length (fl) PrP. In the EVs fractions, apart from a diglycosylated fl-PrP band, a major band at 34 kDa was present which, as judged by the molecular weight (in our system of pre-cast gels, bands run slightly higher than with other systems), could either represent unglycosylated fl-PrP or its diglycosylated C1 fragment ^39^. In order to discriminate between these two forms, we treated the sEVs and the TH fractions with PNGase to enzymatically remove the N-linked glycans (Fig. 3B). This revealed that the dominant band in the EVs fractions is indeed the C1 fragment of PrP (running at 19 kDa upon deglycosylation). To further prove the identity of this N-terminally truncated C1 fragment, membranes were also developed with the POM2 antibody, recognizing the N-terminal half of PrP ^46^. As shown in Suppl. Fig. 1, after PNGase treatment, only unglycosylated fl-PrP (and a faint band corresponding to diglycosylated fl-PrP due to incomplete deglycosylation) was detectable, whereas the fragment corresponding to PrP-C1 was not visible anymore, thus confirming the identity of PrP-C1 in sEVs. When amounts of PrP in sEVs were compared to the corresponding levels in TH (Fig. 3C, D), we demonstrate that total levels of PrP are significantly enriched in brain-derived sEVs (PrP in TH set to 100% ±21.8%; sEVs: 195.5% ±14.9%; *p*= 0.0073). Intriguingly, we found its C1 fragment to be nearly 4-fold increase in sEVs compared to TH (PrP-C1 in TH set to 100% ±13.2%; PrP-C1 in sEVs: 382.7% ±42.5%; *p*= 0.011; Fig. 3C, D). Since the identity of the protease(s) responsible for the α-cleavage of PrP is still unknown ^38, 59^, and in order to rule out the possibility that the enrichment in PrP-C1 is an artefact caused by the collagenase III treatment used for sEV isolation, we also isolated sEVs with collagenase IV. As shown in Suppl. Fig. 2, the PrP-C1 pattern is similar to samples treated with collagenase III.

**Fig. 3.**
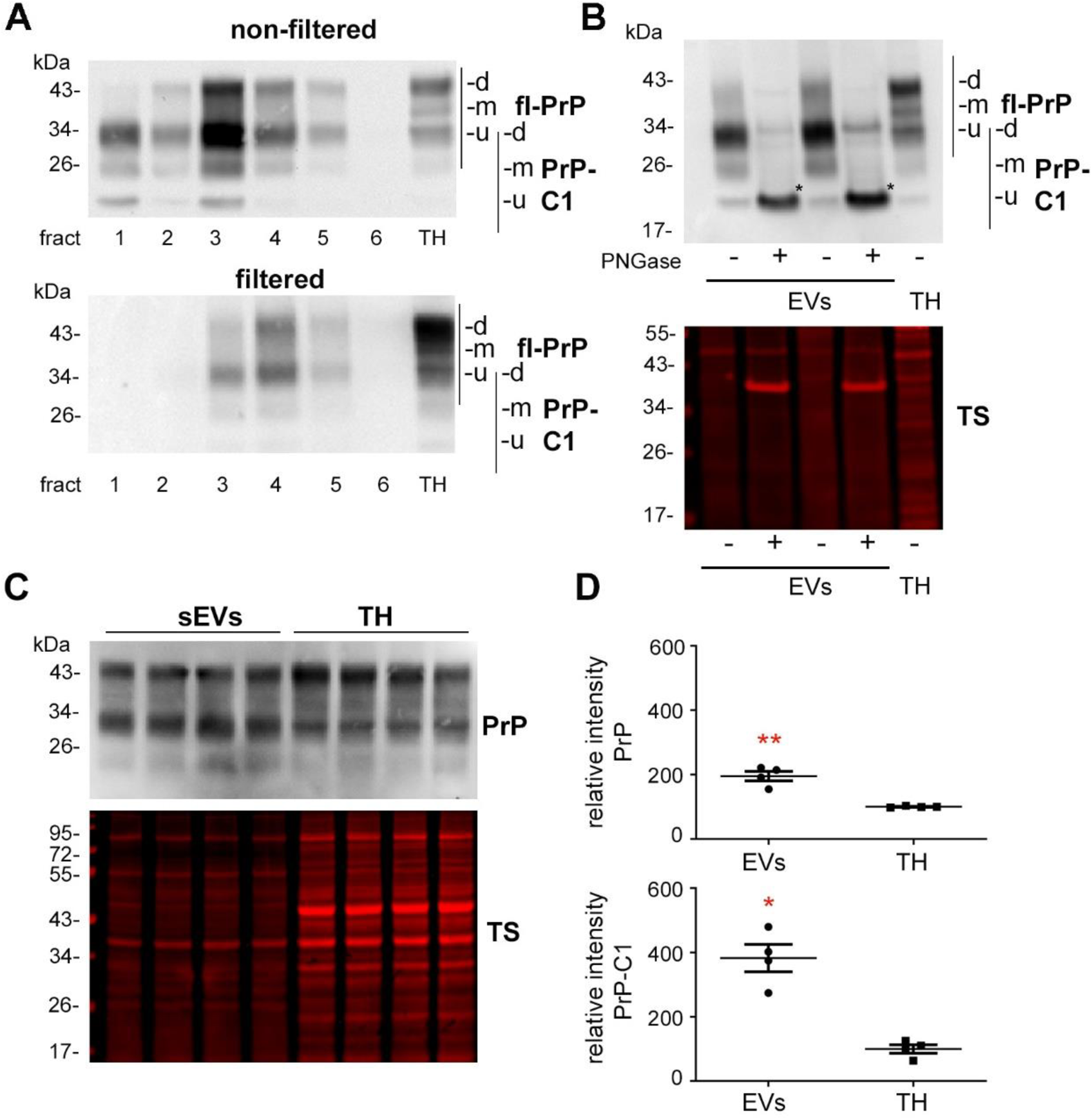
Brain sEVs are enriched in the PrP-C1 fragment. (A) Representative western blots of the six fractions obtained after sucrose gradient centrifugation of non-filtered (EVs) and filtered EVs (sEVs) probed with POM1. A total homogenate (TH) was loaded for comparison purposes. Note that in both cases, PrP in the TH presents a prominent diglycosylated full-length band (fl-PrP) at 43 kDa followed by two other bands, corresponding to mono- and unglycosylated PrP, respectively. In EVs, a major band at 34 kDa is presented which could either correspond to unglycosylated fl-PrP or to its diglycosylated truncated C1 fragment (PrP-C1). (B) Representative western blots of sEVs fractions 3 and 4 treated (+) or not (-) with PNGase F and probed with POM1 and total protein staining (TS). The PNGase assay reveals that the major band at 34 kDa present in the EVs corresponds to the C1 fragment (marked with an asterisk). (C) Representative western blots of pooled sEVs (n=4) compared to their respective total brain homogenates (TH) probed with POM1 and total protein staining (TS). (D) Scatter plots showing quantifications of the comparison between total PrP in the TH vs sEVs, and of PrP-C1 in TH vs sEVs. Each lane was first referred to its total protein staining and then the means were compared. Fl-PrP shows a significant twofold increase in sEVs, whereas PrP-C1 is almost four-fold-increased in sEVs. The mean, SEM and *p* values are reported in the main text.

### Increased release of sEVs by astrocytes and elevated levels of PrP in brain-derived sEVs after tMCAO

To study changes in the relative contribution of different cell populations to the sEV pool in brain at 24 hours after tMCAO, we performed western blot analyses with the brain cell markers described above, comparing sEVs isolated either from shams (n=8) or from tMCAO-operated mice (n=8) (Fig. 4A). We observed that only the levels of the astrocytic marker EAAT1 were significantly increased in sEVs after tMCAO (shams were set to 100% ±5.6% vs tMCAO, 134.2% ±11.1%; *p*= 0.0158), probably indicating that at 24 hours after stroke reperfusion injury, there is an increased release of sEVs by astrocytes. Although there was a slight tendency for oligodendrocytic PLP (shams set to 100% ±4.2% vs tMCAO, 158.4% ±34.1) and microglial P2Y12 (shams set to 100% ±4.3% vs tMCAO, 110.8% ±16.8%) to also be increased in sEVs after tMCAO, this was not significant. The neuronal marker synapsin1 was not changed between shams and tMCAO samples (shams set to 100% ±8.7% vs tMCAO, 102% ±10%).

**Fig. 4.**
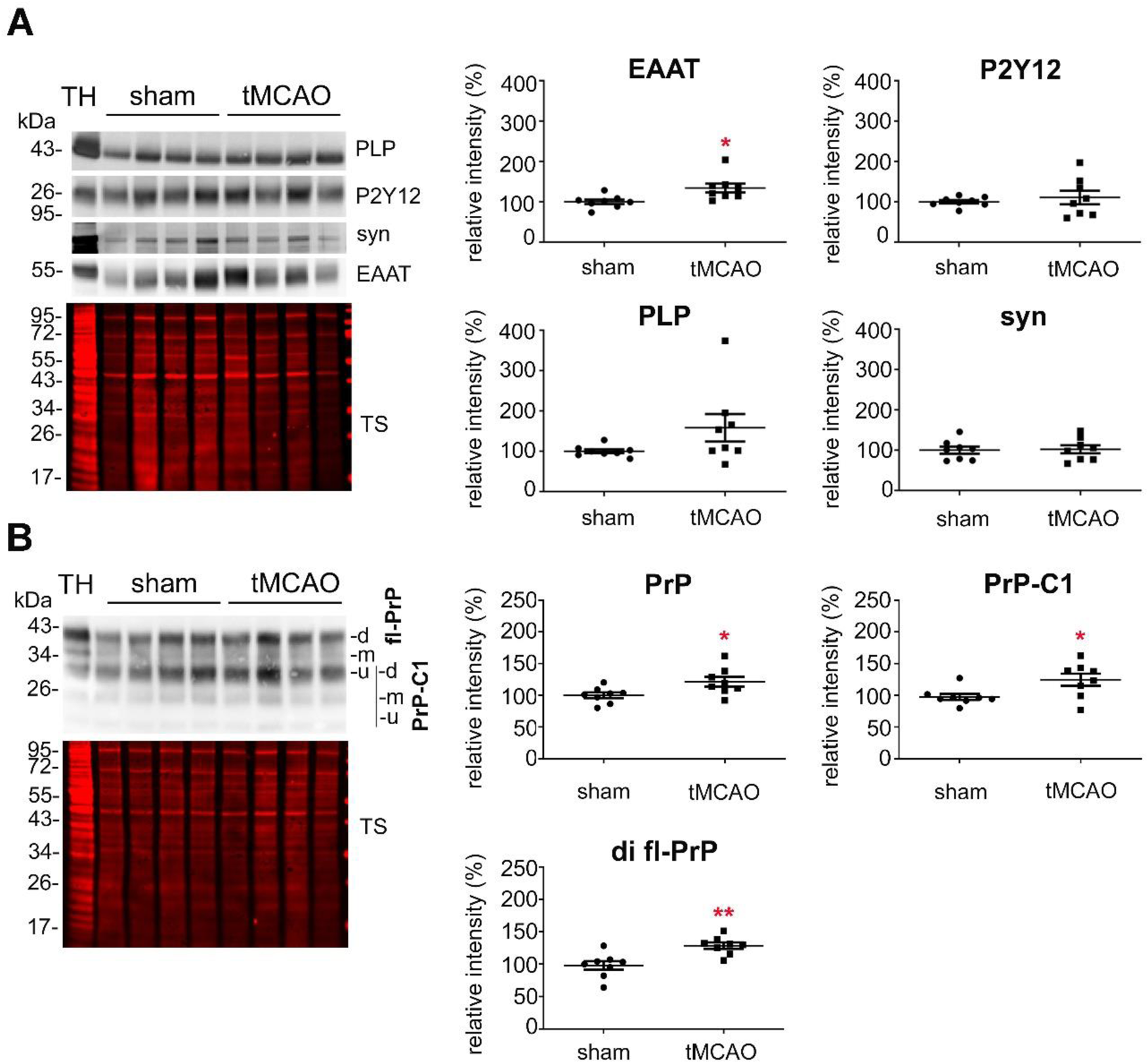
At 24h after tMCAO, astrocytes release higher amounts of sEVs and PrP is increased in sEVs. (A) Representative western blots of sEVs from sham mice (n=8) and mice that underwent the tMCAO procedure (n=8). Blots show the total protein staining (TS) and cell type-specific surface markers: PLP for oligodendrocytes, P2Y12 for microglia, synapsin1 for neurons and EAAT1 for astrocytes. On the right side, scatter plots of relative intensity quantifications, showing a significant increase of the astrocytic marker EAAT1 in tMCAO sEVs compared to sham sEVs. PLP, P2Y12, and synapsin1 were not significantly changed. For quantification each lane was first referred to the TS and then the mean values of the two groups (sham, tMCAO) were compared. (B) Representative western blots of sEVs from sham (n=8) and tMCAO mice (n=8) probed with POM1 antibody and total protein staining (TS). On the right side, scatter plots of relative intensity quantifications showing an increase in total PrP, in PrP-C1 and in the diglycosylated full-length PrP (di fl-PrP) in tMCAO sEVs compared to sham sEVs. The mean, SEM and exact *p* values are reported in the main text.

Notably, a significant increase was also seen for total PrP (Fig. 4B), which was enriched in sEVs from tMCAO samples compared to sham brains (shams set to 100% ±4.4% versus strokes: 121.47.6%; *p*=0.0284). When diglycosylated fl-PrP (di-fl-PrP) and the PrP-C1 bands were quantified individually, we observed that both were significantly increased in sEVs after tMCAO (for PrP-C1: shams set to 100% ±5.2% versus strokes: 124.9% ±9.6%; *p*= 0.039/for di fl-PrP: shams set to 100% ±6.6% versus strokes: 131.3% ±5.2; *p*= 0.0024).

### PrP influences uptake of brain-derived EVs by primary neurons and glia cells

We have demonstrated that PrP in brain-derived EVs is highly enriched in the C1 fragment. This fragment, resulting from the α-cleavage in the middle of the PrP sequence, exposes a hydrophobic domain at its N-terminus ^39^. Both, the structural aspect with a stretch of hydrophobic amino acids and its dependence on proteolytic “activation” are strikingly reminiscent of some viral surface glycoproteins critical for host cell attachment and membrane fusion (for instance those of some paramyxoviruses). In these viruses, the viral envelope fuses with the host cell membrane with the help of two glycoproteins, one that initially attaches the virus and another one that, after an endoproteolytic cleavage, exposes the highly hydrophobic fusion peptide, which integrates into the host cell membrane and mediates the fusion process ^60, 61^. Because of the above-mentioned similarities between PrP-C1 and the fusogenic viral surface proteins, and given the enrichment of PrP-C1 in the EVs membrane, we hypothesized that the C1 fragment might act as a tethering factor and play a role in the uptake of EVs by cells ^39^. To assess the role of PrP in EVs, we incubated primary neuronal cultures from WT mice with labeled sEVs isolated from either WT or PrP-KO mouse brain and fixed them after 1, 3 or 6 hours of incubation. As shown in Fig. 5A, after 1h of incubation, labeled sEVs from WT (shown in white) presented as a rather diffuse staining surrounding the neuronal cell body (marked by phalloidin staining in green), whereas the PrP-KO-sEVs showed a dotty staining at the plasma membrane, and with some of them having already been internalized at this early time point. Surprisingly, we also observed that our primary cultures contained some microglia, despite having been treated with Arabinoside C to eradicate proliferating cells. In several instances, these microglia were found to contain high amounts of PrP-KO-derived sEVs, a feature that we could not observe for WT-derived sEVs (Fig. 5A, Fig 6A). We then established another protocol for primary neuronal cultures from WT mice (low-density culture, LDC), where neurons grow in co-culture with (yet spatially separated from) astrocytes^45^. On the one hand, this approach allows for lower seeding density and, therefore, improved microscopic analysis, while on the other hand, contamination by other brain cell types is sensibly lower. As shown in Fig. 5B, we could again confirm a strong difference in the neuronal uptake behaviour, with PrP-KO-derived sEVs being efficiently taken up in contrast to sEVs from WT brain (which at 1h of incubation were hardly detectable in the culture). Moreover, although microglia were much less and therefore more difficult to find in this type of culture, the few microglia cells that could be identified were also highly decorated with PrP-KO-sEVs, whereas microglia showed less positive EV-associated labelling when treated with WT-derived sEVs. The quantification in Fig. 5C shows that the neuronal uptake in the primary culture with higher amounts of microglia (HDC) was almost twofold increased for the PrP-KO-derived sEVs than for WT-derived sEVs (WT-sEVs: set to 100% ±5.3%; PrP-KO-sEVs: 185.8% ±12.5%; *p*≤0.0001), whereas in the LDC, with low microglia content, neuronal uptake of sEVs from PrP-KO brain was about three times higher compared to WT-sEVs (WT-sEVs: set to 100% ±8.6%; PrP-KO-sEVs: 323.3% ±22.1%; *p*≤0.0001). We hypothesize that this difference between the culture conditions could either be a consequence of the lower number of neurons or the reduction in microglia in the LDC, both resulting in a higher sEV-to-neuron ratio. In order to confirm that microglia were taking up high amounts of PrP-KO-sEVs, we established a mixed glial cell cultures (mainly containing microglia and astrocytes) and incubated them with either WT-sEVs or PrP-KO-sEVs for 3h. This type of culture allowed us to measure the sEVs uptake not only by microglia but also by astrocytes (which were absent in our previous primary neuronal cultures). After incubation with sEVs we labelled the two populations with cell specific markers (GLAST-1 for astrocytes and CD11b for microglia) and performed flow cytometric analysis. As shown in Fig. 6B, we could confirm that microglia take up more PrP-KO-sEVs than WT-sEVs (WT-sEVs set to 100% ±2.5% vs PrP-KO-sEVs: 163.2% ±22.3%; *p*=0.035). Therefore, as observed for neurons and microglia at 1h, PrP-KO-sEVS were also taken up more efficiently by astrocytes (WT-sEVs set to 100% ±2.6% vs PrP-KO-sEVs: 166.9% ±16.6%; *p*=0.0068).

**Fig. 5.**
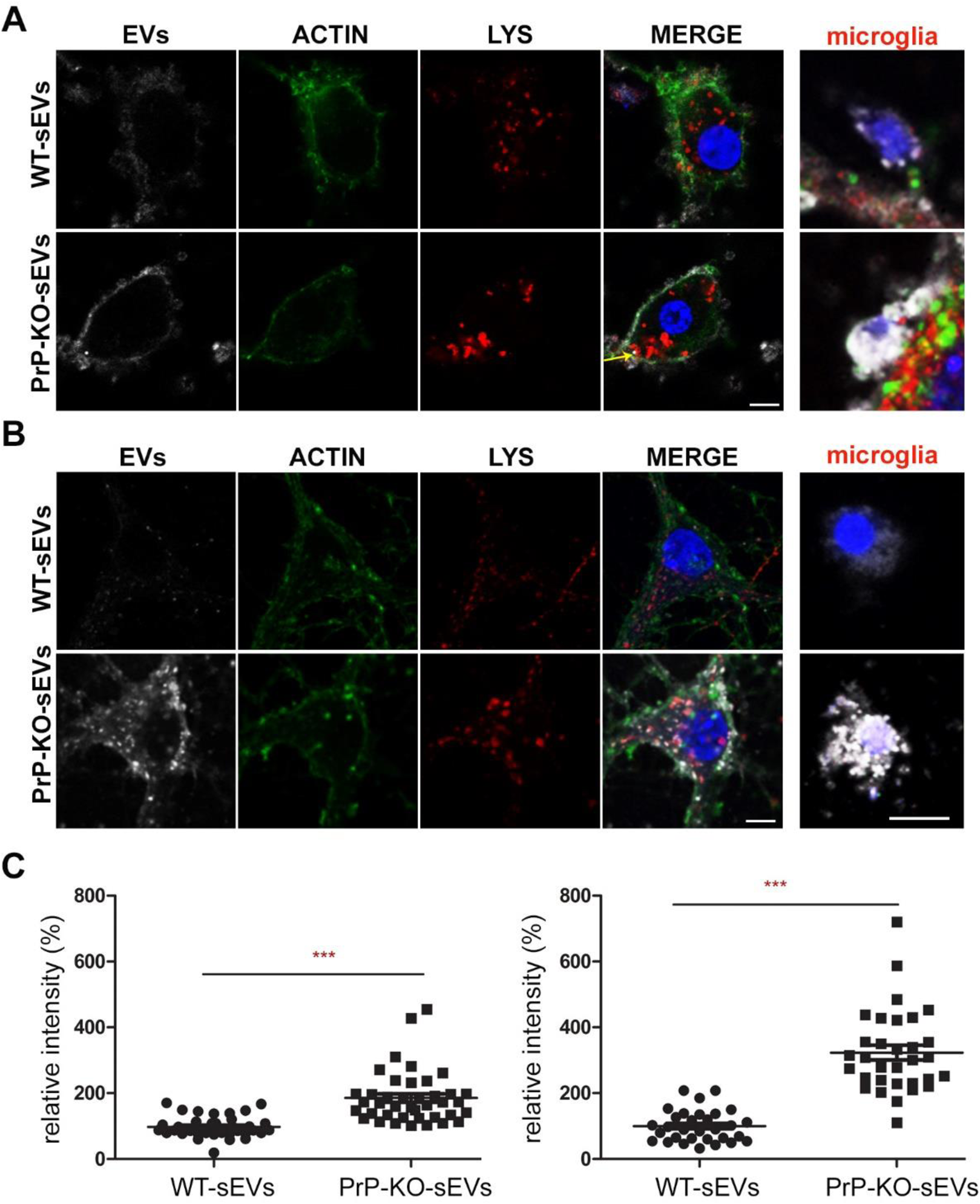
PrP influences brain sEVs uptake by primary neurons. (A) Representative confocal microscopy images of primary neurons from WT mice in high-density culture incubated for 1h with sEVs isolated from WT mouse brains (WT-sEVs) or from *Prnp*^0/0^ mouse brains (PrP-KO-sEVs) labelled with mCLING dye. sEVs signals are shown in white. Neurons were stained with phalloidin to visualize F-actin (green), the lysosomal marker LAMP-1 (red), and with DAPI (blue; to visualize the nucleus). Note that after 1h of incubation, WT-sEVs present with diffuse staining surrounding the neuronal plasma membrane, whereas PrP-KO-sEVs show with a dotty staining at the neuronal plasma membrane, but some PrP-KO-sEVs are also found inside the neuronal body (yellow arrow). In this high-density culture (HDC), other cell types (seemingly microglia) were observed to take up a few WT-sEVs, yet conspicuously higher amounts of PrP-KO-sEVs. (B) Representative confocal images of low-density culture (LDC) primary neurons from WT mice incubated for 1h with sEVs isolated from WT (WT-sEVs) or PrP-KO mouse brains (PrP-KO-sEVs) labelled with mCLING as in (A). Here again, PrP-KO-sEVs are taken up by neurons more readily than WT-sEVs and what we presumed to be microglia cells (see Fig. 6) showed the same engulfment pattern of sEVs as in (A). Scale bar is 5 µm. (C) Scatter plot showing intensity of sEVs quantification in high-density culture (HDC, on the left) and low-density primary neuronal cultures (LDC, on the right). PrP-KO-sEVs are significantly more taken up by neurons than WT-sEVs. The mean, SEM and exact *p* values are given in the main text.

**Fig. 6.**
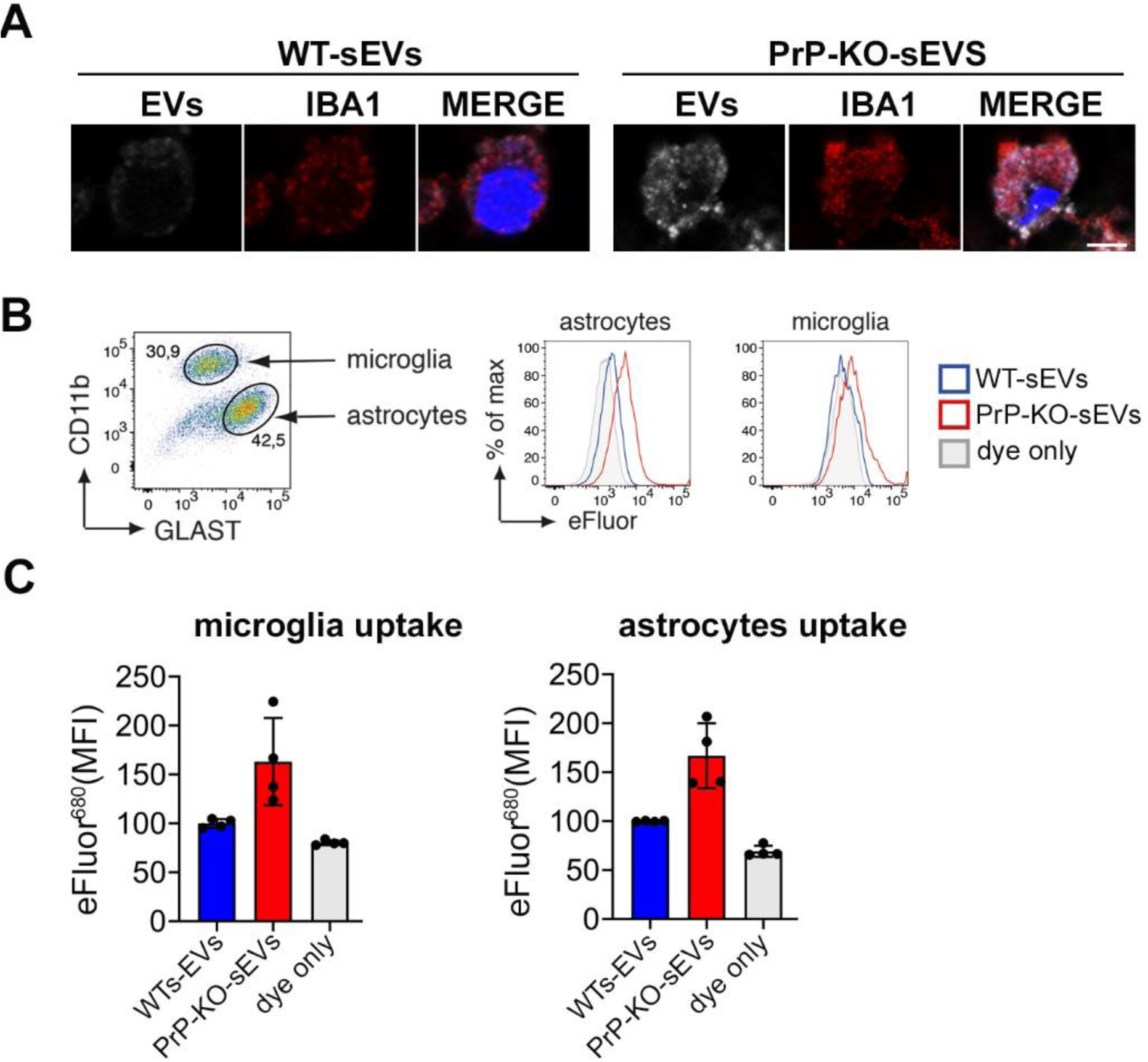
PrP influences the uptake of brain-derived sEVs by microglia and astrocytes. (A) Confocal microscopy images showing that, what we observed as non-neuronal cells in our cultures, are indeed microglia, as they show staining for IBA1. As in Fig. 5, sEVs are labelled with mCLING and shown in white. DAPI (in blue) is used as a nuclear staining. (B) Representative FACS plot of astrocytes (GLAST+) and microglia (CD11b+) from mixed cultured glia cells (n=4). Mixed glia cells from WT mice were incubated for 3h with either WT-sEVs or PrP-KO-sEVs labelled with mCLING and were analysed for sEVs uptake using flow cytometry. On the right side, histograms showing the intensity of the sEVs fluorescence measured either from astrocytes or microglia. Note that in both cases the intensity is higher for PrP-KO-sEVs (C) Bar scatter plot of normalized fluorescence intensity show that both, microglia and astrocytes, take up PrP-KO-sEVs more efficiently than WT-sEVs. The exact means, SEM and *p* values are reported in the main text.

Regarding the neuronal uptake at 3h (Fig. 7A), treatment with WT-sEVs in the HDC again revealed a diffuse staining of the sEV-labeling at the plasma membrane, although seemingly more intense than after 1h (Fig. 5A). In addition, some sEVs could now be detected as vesicle-like structures at the plasma membrane or even internalized and colocalizing with the lysosomal marker LAMP-1, lysosomes being described as one of the major targets of internalized EVs ^62, 63^. In the case of PrP-KO-sEVs, although some signal was still detected at the plasma membrane, the majority at this time point was observed inside the neurons and colocalizing with LAMP-1. Quantification of sEVs signal intensity inside neurons (Fig. 7C) showed that at 3h still about the double amount of PrP-KO-sEVs was taken up compared to WT-sEVs (WT-sEVs set to 100% ±7.45% vs PrP-KO-sEVs: 219% ±16.3%; *p*≤0.0001).

**Fig. 7.**
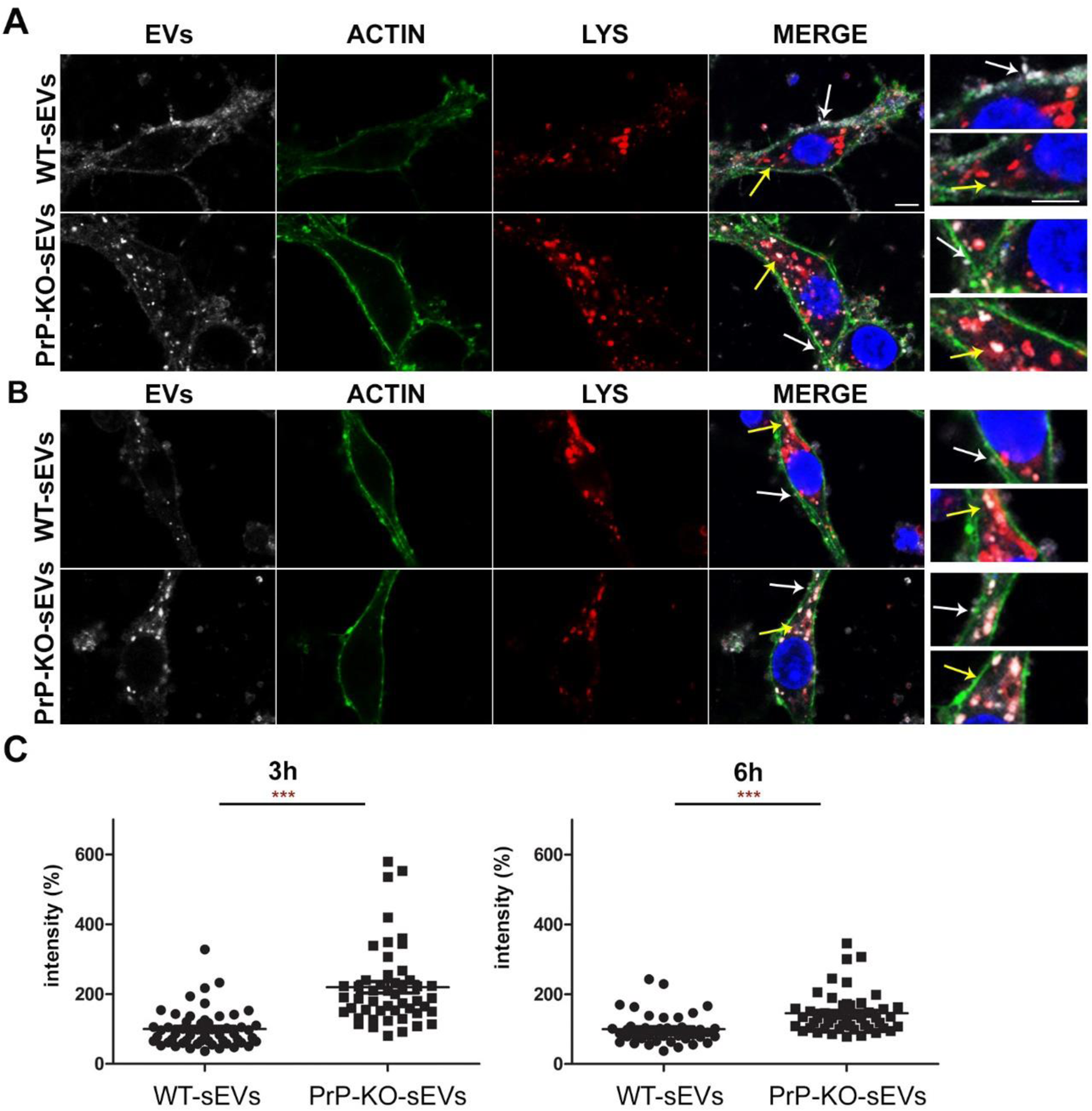
Brain-derived sEVs are found in lysosomes after 3 and 6h of incubation. (A) Representative confocal microscopy images of primary neurons from WT mice (labelled with phalloidin, in green) in high-density culture (HDC) incubated for 3h with WT-sEVs or PrP-KO-sEVs labelled with mCLING (white signal). The lysosomal marker LAMP-1 is shown in red and nuclei are stained with DAPI (in blue). Note that, after 3h, both WT- and PrP-KO-sEVs are co-localizing with LAMP-1, although we consistently observed a much higher level of colocalization for PrP-KO-sEVs. Instead, a diffuse mCLING staining at the membrane is found for WT-sEVs. White arrows show sEVs that are at the plasma membrane (PM) whereas yellow arrows indicate colocalization of sEVs with LAMP-1. Note that for WT-sEVs there are many sEVS at the PM whereas for PrP-KO-sEVs there are much more sEVs colocalizing with LAMP-1.(B) Representative confocal microscopy images showing primary neurons as in (A) but incubated with sEVs for 6h. sEV identity and stainings as in (A). Note that after 6h sEV-associated signals are mainly found in colocalization with lysosomes. Scale bar is 5 µm. (C) Scatter plots showing intensity quantifications of WT-sEVs and PrP-KOsEVs in primary neurons at 3h (on the left) and 6h (on the right) of incubation in the high-density cultures, HDC. As for 1h (shown in Fig. 5), PrP-KO-sEVs are significantly more taken by neurons than WT-sEVs. Mean, SEM and exact *p* values are reported in the main text.

At the latest time point assessed in this study (6h of incubation), the vast majority of both, WT- and PrP-KO-sEVs, were found inside neurons and colocalizing with LAMP-1 (Fig. 7B). Again, quantification of the sEVs-associated fluorescence signal present inside the neurons still revealed a higher amount of uptaken sEVs from PrP-KO compared to those derived from WT brain (WT-sEVs set to 100% ±6.6% vs PrP-KO-sEVs: 145.37% ±8.56%; *p*≤0.0001). For the LDC system (Suppl. Fig. 3), results obtained at 3h and 6h were similar to the ones mentioned above(for 3h incubation: WT-sEVs set to 100% ±3.46% vs PrP-KO-sEVs: 214% ±17.58%; *p*≤0.0001/for 6h incubation: WT-sEVs set to 100% ±5.6% vs PrP-KO-sEVs: 187% ±14%; *p*≤0.0001). In contrast to the 1h incubation (Fig. 5), the absence of differences between the two culture systems at these time points can probably be explained by the fact that there were no more free sEVs present in the media.

## DISCUSSION

In the present study, we have characterized sEVs in the murine brain in steady-state conditions and after 24h of stroke reperfusion. We show that, under physiological conditions, microglia are the main sEVs source in brain. This situation changes at 24h after induced stroke-reperfusion, where a significant increase of astrocytic sEVs was observed. Furthermore, we show that brain-derived sEVs are enriched in PrP, and particularly in its C1 fragment, with consequences in the regulation of vesicle uptake by recipient cells. Thus, when sEVs lack PrP, their uptake by neurons is increased and, conspicuously, they are also more readily taken up by microglia and astrocytes. We also demonstrate that, 24 hours after stroke, the amount of PrP (and PrP-C1) in the brain sEVs pool is significantly increased, which may have functional consequences in intercellular communication after stroke (a graphical summary of the principal findings is depicted in Fig. 8).

**Fig. 8.**
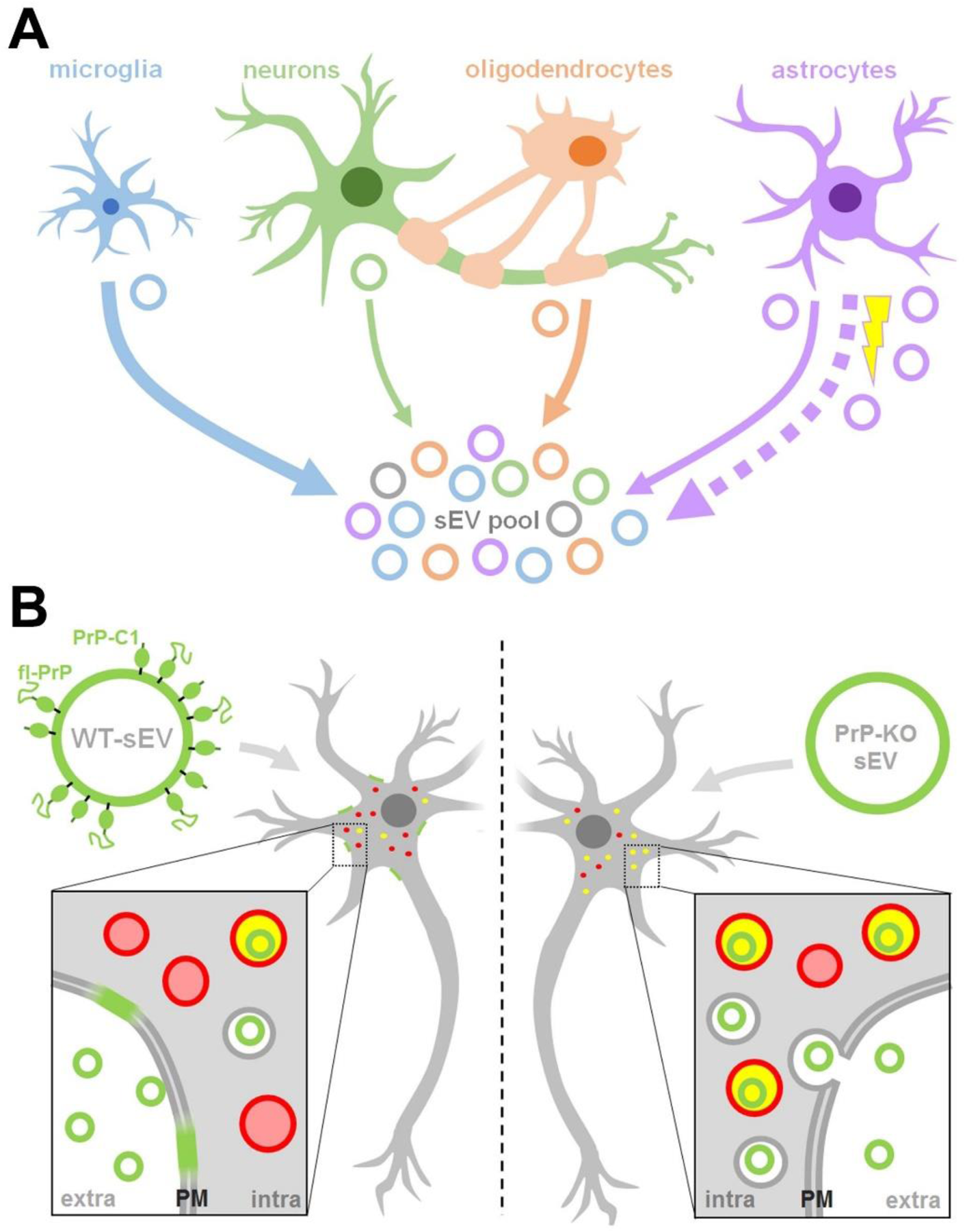
Summarizing scheme. (A) Schematic representation of the relative contribution of different brain cell types to the sEV pool in murine brain. Under physiological conditions, microglia appear to be the main contributor to the sEV pool, followed by oligodendrocytes and astrocytes, while neurons contribute relatively little (contribution indicated by thickness of solid arrows). Upon experimental stroke (indicated by the yellow thunderbolt), astrocytic release of sEVs is upregulated and astrocytes appear to be the main contributor at 24 hours after reperfusion (indicated by the bold dotted arrow). sEVs are depicted as circles and the color refers to their cellular origin. Note that a fraction of sEVs (grey) may also be released by other cell types not assessed here, such as pericytes or endothelial cells. (B) Differential uptake of WT-sEVs and PrP-KO-sEVs may be influenced by PrP. WT-sEVs (on the left) are packed with fl-PrP and its truncated C1 fragment ending with a stretch of hydrophobic amino acids. These sEVs are relatively slowly taken up by neurons and rather seem to fuse with the plasma membrane (PM). In contrast, sEVs lacking PrP (PrP-KO-sEV; on the right) are rapidly endocytosed and transported to lysosomes (red circles). Colocalization (i.e. lysosomes containing sEVs) is indicated in yellow. Similar observations have been made with microglia or astrocytes as recipient cells. Note that other surface proteins and cargo of sEVs, as well as the lipid bilayer of vesicles, lysosomes, and sEVs, are not depicted here to simplify matters.

EVs have become an intense field of research, not only because they represent a novel form of cell-to-cell communication able to bridge wide distances, but also by their potential applicability as therapeutic tools. In the case of brain disorders, they are very attractive because of their property to cross the BBB when genetically modified to target the brain ^64^. A protective role of EVs derived from mesenchymal cells in a variety of brain insults and disorders ^65^, including stroke ^66–69^, has also been shown. Until recently, most of the EVs assessed in various studies were isolated from cell culture supernatants or body fluids, but the isolation of EVs from complex tissue such as the brain parenchyma has been increasingly reported only in the last few years ^42, 70, 71^.

Despite enormous progress in recent years, the isolation and characterization of EVs are still a challenge, which is mainly due to their size. One of the aims of our study was to characterize the differential contribution of the brain cell populations to the EVs pool in normal conditions and to assess potential changes after 24 hours of ischemia-reperfusion in the tMCAO mouse model of stroke. We modified a protocol of EVs isolation from brain ^42^ by including a filtration step in order to differentiate between small EVs (≤200 nm) and medium/large EVs (≥ 200nm). We performed this step to yield a homogeneous population for our downstream analyses. As assessed by EM and NTA, we could observe that, without this additional filtration step, there is a heterogeneous population of brain EVs. Based on size measurements and the presence of Alix, a resident protein of MVBs, we think that our brain-derived sEVs represent a mixture of bona fide exosomes and small microvesicles ^48^.

Intuitively, it may seem reasonable that exosomes (released upon fusion of MVBs with the plasma membrane) and microvesicles (generated by “budding” from the plasma membrane) are released upon different stimuli and that some cell types are probably more prone to release one type over the other. However, it is not known whether this scenario is really the case.

Because the cargo of EVs is cell type-dependent and includes proteins that reflect their cellular origin, we sought to investigate the relative contribution of different cell types to the brain sEVs pool by assessing the relative enrichment (compared to TH) of certain brain cell markers by western blot. We show that, in steady-state conditions, P2Y12 and TMEM-119, two known markers of microglia, are significantly enriched in the pool of sEVs of mouse brain. On the contrary, the neuronal markers synapsin and SNAP-25 showed a significant relative decrease, indicating a poor contribution to the total sEVs brain pool. Oligodendrocytes (labelled with PLP and CNP1) and astrocytes (with EAAT-1 and EAAT-2 as exclusive markers), showed no enrichment in sEVs when compared to the TH, and therefore, we concluded they are not the major populations that contribute to the brain sEVs pool in steady-state conditions. We cannot rule out the possibility that certain proteins are more heavily packed into EVs than others, but the use of two markers for each cell type and the fact they were behaving similarly, reduces the risk of misinterpretation and ensures more reliable results regarding the relative contribution of different cells. Relative quantification of EVs in brain is not an easy issue. In a recent paper, Silverman *et al.* ^72^ have also characterized the relative proportion of EVs derived from different brain cell populations. By means of flow cytometry and in contrast to our findings, they observed an increased proportion of astrocytic and neuronal markers in both, brain and in spinal cord samples. The discrepancy with our results could be a consequence of (i) investigation of different subpopulations of EVs (note that we included a 0.2 µm filter step) and (ii) the detection limit of the flow cytometry possibly excluding some small vesicles that were considered as debris (which are included in the western blot analyses of our study). It has been calculated that sEVs are at least one order of magnitude more frequent than medium/large EVs ^73, 74^ and this pool may have been lost by flow cytometry measurements.

What we observed under the same settings is that, 24 after stroke reperfusion, brain sEV landscape was changing as sEVs containing the astrocytic marker EAAT-1 were significantly increased. Astrocytes have a myriad of important functions in the brain, ranging from regulation of synaptic transmission, modulation of neuronal excitability, BBB formation, and regulation of blood flow. Since they are relatively resistant to glucose and oxygen deprivation, they cope better than neurons with ischemic insults and, at present, they are raising attention as potential targets in stroke therapy ^75^. At earlier times points after ischemia, astrocytes proliferate, become hypertrophic (reactive astrogliosis) and, after a few days, start to form the glial scar with both, potential benefits but also detrimental effects ^76^. It has been shown that astrocytes start proliferating from one day after injury ^77^. Likewise, microglia are significantly increased at 24h in a mouse model of stroke ^19^. Since there is close communication between astrocytes and microglia after injury, and given that activated astrocytes can in turn contribute to the activation of distant microglia after ischemia ^78^, it may be that the release of EVs from astrocytes at 24h is a factor facilitating the spread of reactive microglia. Moreover, astrocytes present neuroprotective features after stroke ^79^ and neurons exposed to reactive oxygen species (ROS) showed increased survival when treated with astrocytic EVs ^80^. Whether the relative increase in astrocytic sEVs is related to microglia proliferation, the formation of the glial scar, or for protection against ROS, clearly deserves further studies. Interestingly, Guitart *et al*. ^36^ showed that exosomes released by astrocytes under hypoxic conditions (using the *in vitro* model of oxygen-glucose deprivation, OGD) conferred protection to neurons. Of note, this activity was dependent on the presence of PrP, which they found increased in exosomes derived from stressed astrocytes. Contrarily, exosomes from stressed astrocytes devoid of PrP were not protective. In our experiments, apart from the increase of sEVs released by astrocytes, we also observed an increase of sEV-associated PrP levels after stroke. Due to technical limitations, we could not assess whether this upregulation is caused by the relative increase of astrocytic sEVs. However, in view of the reported neuroprotective effects of PrP after ischemia ^30, 35 Doeppner, 2015#933^, this increase in both, astrocytic sEVs and sEV-associated PrP amounts, could represent a protective feedback mechanism to counteract the oxidative stress present in the first hours after stroke. Although an increase in PrP levels has been observed in animal models of stroke and in humans after ischemic insult ^33, 81–83^, the present study is the first to report elevated levels of this protein in brain-derived sEVs. However, further studies are clearly required to elucidate its consequences.

We also demonstrate that, in steady-state, the proteolytically generated PrP-C1 fragment is particularly enriched in brain sEVs. Vella *et al*. ^84^ already found that PrP present in EVs could not be recognized with antibodies against the N-terminal part of the protein, indicating truncation. In support of that study, we show here that the main PrP forms packed onto sEVs in the brain are PrP-C1 together with fl-PrP (with both largely being diglycosylated). We had already suggested a parallelism between the hydrophobic domain, which is N-terminally exposed in C1 after the α-cleavage of PrP, and the fusion peptide of certain viral surface proteins ^39^. Because the latter allows viruses to dock to and fuse with the host cells, we hypothesized that PrP-C1 on EVs could perform a similar function, such as tethering EVs to recipient cells and/or facilitating their uptake. The fact that EVs also share many similarities with viruses ^85, 86^ would reinforce our hypothesis. Surprisingly, we found that, as early as one hour of incubation, brain sEVs lacking PrP are readily taken up and present inside neurons (and glia cells), whereas sEVs from WT mice are not taken up at this time-point, but are rather present as diffuse staining surrounding the neuronal body. At later time points, although WT-sEVs are taken up and start to colocalize with lysosomes, there are still significant differences compared with PrP-KO sEVs, with the latter being massively taken up and colocalizing with lysosomes ^87^. This seems counterintuitive with our initial hypothesis since our findings rather indicate that a lack of PrP leads to more efficient uptake. However, it has been shown that EVs can either fuse with the plasma membrane, thereby releasing their content into the cytoplasm, or they can be taken up by clathrin/caveolin-dependent mechanisms or by phagocytosis and macropinocytosis ^88^. Since we always incubated the same amount of sEVs but consistently detected more endocytosed PrP-KO sEVs, this could indicate that presence of PrP and/or PrP-C1 (in WT-sEVs) indeed influences fusions events. This may also be supported by the diffuse sEV-associated staining at the plasma membrane found at 1h and 3h for WT-sEVs. In contrast, in the absence of PrP, it seems that vesicles are taken up more quickly by other mechanisms and delivered to organelles (i.e. lysosomes). Thus, modulation of PrP composition could regulate EV fusion versus uptake and, consequently, influence whether EV cargo is released to the cytoplasm or intracellular compartments, respectively. It has been reported that fusion between EVs and plasma membrane in a cell-free system requires the presence of proteins at the surface of both entities and an acidic pH ^89^. Given that, at a later time-point, we observed both WT- and PrP-KO-sEVs to colocalize with LAMP-1, final lysosomal targeting of sEVs seems to be PrP-independent. In general, intracellular cargo delivery of EVs is still poorly understood and further studies, as for the potential role of PrP in these processes, are clearly needed to clarify this important point in EV biology.

Interestingly, the uptake of sEVs lacking PrP by microglia and by astrocytes was likewise highly increased. This result could point out to a role of PrP in EV recognition by the immune system. It is striking that PrP is highly abundant in organs that possess immunological privilege, such as brain, placenta, and testicles ^90^. Fittingly, in several inflammatory processes, such as ischemia, brain trauma, and EAE, the inflammatory damage is exacerbated when PrP is absent, suggesting a role in immune quiescence ^90^.

In conclusion, the present study describes that microglia are the main contributors to the sEV pool in brain under physiological conditions and that brain sEVs are enriched in PrP-C1, which may modulate EV uptake in recipient cells. After stroke, astrocytes increasingly contribute to the sEV pool and elevated levels of sEV-associated PrP are detected. These findings add novel insight to previous studies indicating a major role of both, PrP and EVs in the brain under physiological and ischemic conditions.

## ACKNOWLEDGMENTS

The authors would like to thank Oliver Schnapauff and Anika Ruhl for performing the animal model of tMCAO. This work was supported by the “Werner Otto Stiftung” (WOS) (Grant 13/91 to BP) and by the “Hermann und Lily Schilling Stiftung” (Grant to TM), the German Research Council SFB 1328 (project A13, grant to TM), GRK1459 (to BM and MG), and SFB877 (project A12 to MG). Work of HCA is supported by the WOS and the CJD Foundation, Inc.

