## Supplementary material for "Brain-Derived Extracellular Vesicles are Highly Enriched in the Prion Protein and Its C1 Fragment: Relevance for Cellular Uptake and Implications in Stroke": Suppl. Fig 1, Suppl. Fig 2 and Suppl.3

**Suppl. Fig. 1.- Brain-derived sEVs treated with PNGase**. Representative western blot of sEVs and TH treated (+) or not (-) with PNGase probed with POM2 antibody (recognising the N-terminal part of PrP [46]) and subsequently re-probed with POM1 antibody (directed against the C-terminal part of PrP). Note that POM2 is unable to recognise the PrP-C1 fragment after PNGase treatment, further proving the identity of this band. "TS" is total protein staining used as a loading control.

**Suppl. Fig. 2.- Brain-sEVs extracted with collagenase IV show the same PrP-C1 pattern.** Representative western blot of the six fractions of sEVs isolated using collagenase IV obtained after sucrose gradient centrifugation. Total protein staining (TS) is shown as a loading control. The Golgi resident protein GM130 is used as a non-sEV marker while Alix and Flotilin-1 serve as sEV markers. Blots probed with POM1 show that the C1-to-PrP pattern is conserved and that it is not an artefact caused by the sEV isolation procedure.

**Suppl. Fig. 3.- Low-density primary neuronal cultures (LDC) also show higher uptake of brain-derived PrP-KO-sEVs.** (A) Representative confocal microscopy images showing that low-density primary neurons (co-cultured with an astrocyte feeder layer until start of the sEV treatment) incubated for 3h with mCLING-labelled WT-sEVS or PrP-KO-sEVs. PrP-KO-sEVs are more readily taken up than WT-sEVs (as shown before in the high-density neuronal culture, HDCFig. 5). Labelled in white are sEVs, neurons are stained with phalloidin (green), the lysosomal marker LAMP-1 (red) and DAPI (in blue). (B) Representative confocal microscopy images of low-density primary neurons incubated for 6h with either WT-sEVs or PrP-KO-sEVs;as in A). In here, sEVs derived from both, WT and PrP-KO brains, are colocalizing with lysosomes, yet amounts taken up are significantly higher for PrP-KO-sEVs as shown in the scatter plot of the quantification (in C). Scale bar is 5 µm. Mean, SEM and exact *p* values are reported in the main text.

SUPPL. FIG. 1

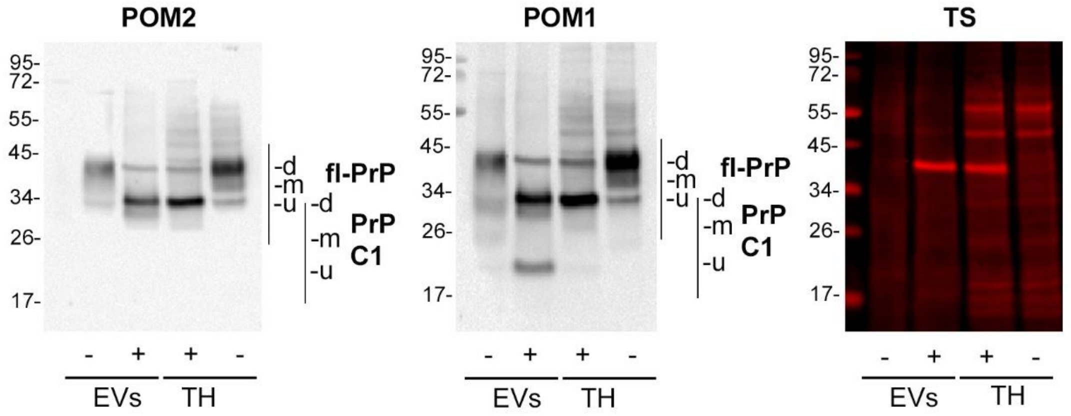


SUPPL. FIG 2

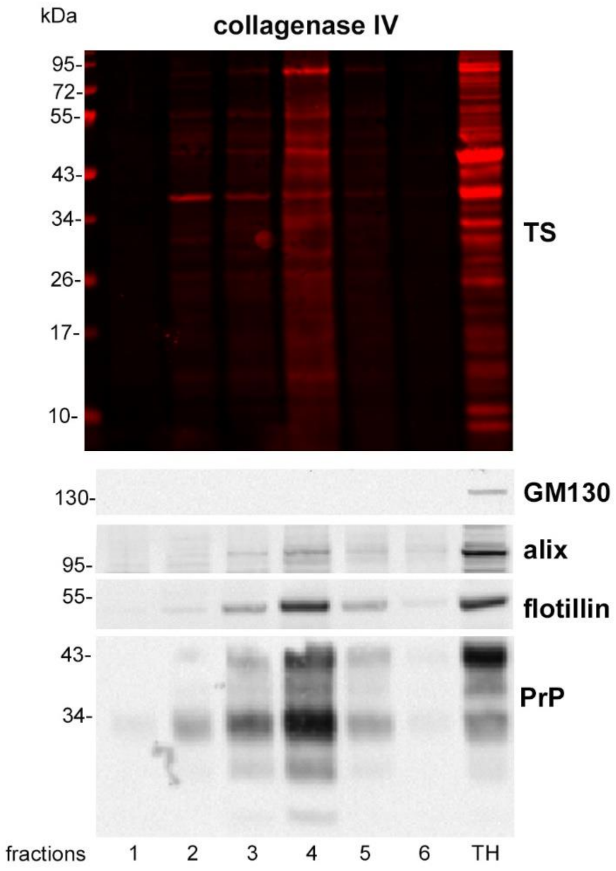

SUPPL.FIG.3

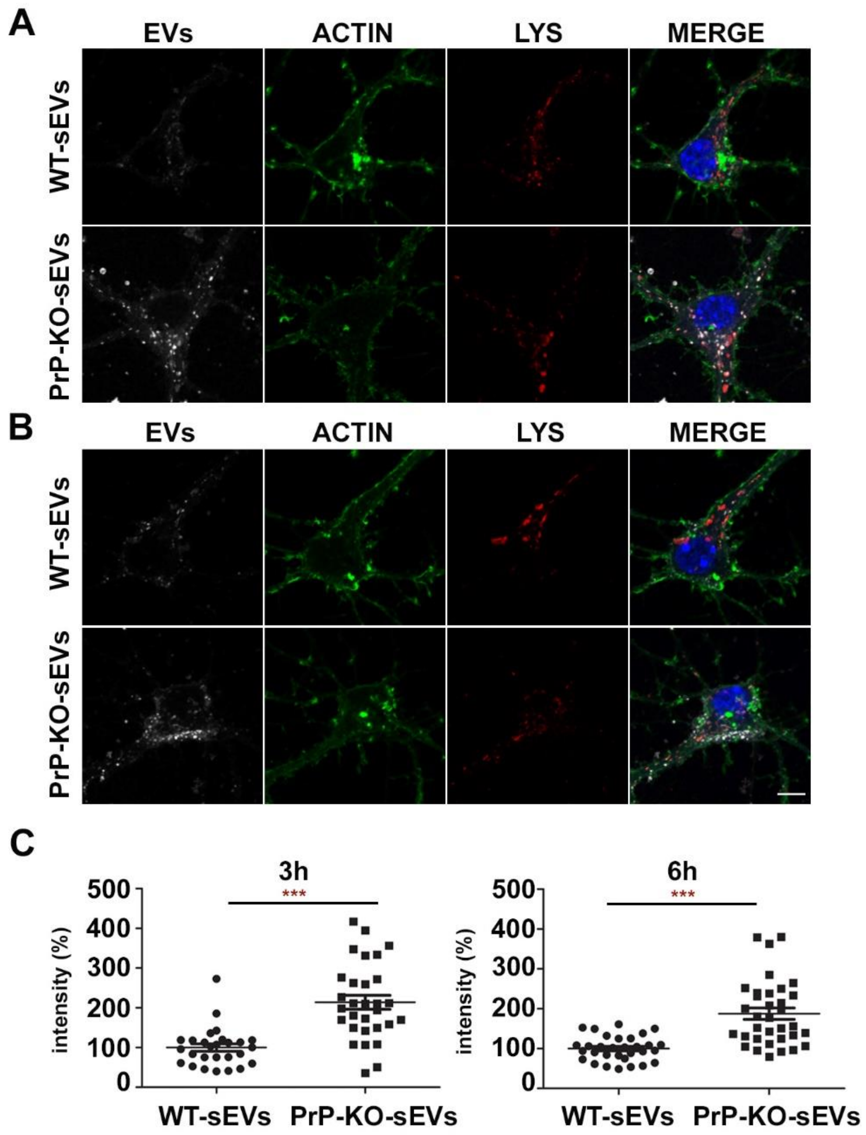

A
EVs
ACTIN
LYS
MERGE
WT-sEVs
PrP-KO-sEVs
B
EVs
ACTIN
LYS
MERGE
WT-sEVs
PrP-KO-sEVs
C
3h
***
intensity (%)
500
400
300
200
100
0
WT-sEVs
PrP-KO-sEVs
6h
***
intensity (%)
500
400
300
200
100
0
WT-sEVs
PrP-KO-sEVs
